# Casual Versus Fractal Modularity in Mammalian Social Organisation

**DOI:** 10.1101/2025.11.26.690800

**Authors:** R.I.M. Dunbar

## Abstract

Multilevel social systems in birds and mammals have attracted considerable interest, not least because they share some similarities with the natural structural organisation of human societies. I use a fractal analysis of a taxonomically wide range of species reported to have multilevel social systems to show that these in fact involve two very different phenomena: those that are fractally modular and those that are casually modular. The first have a strictly fractal structure identical to that found in human societies; the second lack formal structural coherence and are closer in form to casual herds. The former are associated with larger neocortices than the latter, suggesting that they may be cognitively more demanding. Understanding the evolution of multilevel systems and the cognition that underpins them obliges us to adopt a more nuanced approach to social evolution.

## Introduction

There has been considerable recent interest in the fact that some species of group-living mammals have multilevel societies in which small groups are bound together into progressively larger groupings (see, for example, Grueter et al. 2012; Camerlenghi & Papageorgiou 2025). Examples include elephants and orcas (Hill et al. 2008), humpback and sperm whales (Cantor et al. 2015; Whitehead 2024), spotted hyaena (Dunbar 2024), snubnosed (*Rhinopithecus*) (Bleisch et al. 1993; Bennett & Davies 1994; Kirkpatrick 1996) and proboscis (*Nasalis*) (Yaeger 1990, 1991) monkeys, hamadryas (*Papio hamadryas*) (Kummer 1968; Hill et al. 2008) and Guinea (*P. papio*) baboons (Dunbar & Nathan 1972; Fischer et al. 2017), drills and mandrills (*Mandrillus* spp.) (Bret et al. 2013), and gelada (*Theropithecus*) (Hill et al. 2008; MacCarron & Dunbar 2016). Plausible claims for multilevel sociality have also been made for birds. Examples have included bell miners (Painter et al. 2000), Guinea fowl (Papageorgiou et al. 2019) and fairy wrens (Camerlenghi et al. 2022).

Among the primates, *Colobus angolensis* has attracted particular interest because at least two of its East African populations (Nyungwe in Rwanda: Fimbel et al. 2001; Miller 2019; Ruwenzori in southwestern Uganda: Stead & Treichot 2019) are reported to form groups in excess of 300 individuals that are an order of magnitude larger than this species (or any other *Colobus* species) forms anywhere else in Africa. A bimodal distribution of group sizes of this kind has been interpreted as *prima facie* evidence for a multilevel structure (Dunbar et al. 2018; Dunbar 2024).

Multilevel social organisation is also characteristic of human societies. Small scale human ethnographic societies exhibit a very consistent modular structure with a scaling ratio (the ratio of average group sizes between successive layers) of ∼3 (Hill & Dunbar 2003; Zhou et al. 2005; Hamilton et al. 2007; Sandeford 2018; Bird et al. 2019; Dunbar 2020). In humans, this multilevel layering characterises both the organisational structure of communities as well as egocentric personal social networks (Dunbar 2020). Similar internal structuring of social relationships within groups, with the same scaling ratios and the same group sizes as found in humans, has been noted in chimpanzees (Escribano et al. 2022), with some evidence to suggest that this may also be true of a number of other primate and non-primate species (Hill et al. 2008; Dunbar 2024).

Although the social arrangements of all these species share a degree of similarity in that small groups come together to form larger groups, an important criterion is whether these higher level groupings have a significant degree of demographic consistency (supergroups are composed only of specific subgroups). Hill et al. (2008) showed that multilevel structures of the first kind have a distinctive fractal structure in which the successive layers have a consistent pattern with each layer being approximately three times the size of the layer beneath it, as is the case in human social networks (Zhou et al. 2005; Hamilton et al. 2007; Sandeford 2018; Dunbar 2020). In other cases, supergroups seem to be more casual in their formation and structure, with large herds forming and disbanding as a function of local ecological conditions in a manner similar to ungulate herds (e.g. many colobines). When the formation of larger scale groupings is more casual, it is likely that very different cognitive demands are involved (Dunbar & Shultz 2021a). This is because holding bonded groups together in the face of the very strong centrifugal forces that drive animals apart is cognitively demanding, and becomes increasingly so as group size increases (Dunbar 2025). Treating these two kinds of multilevel groupings as synonymous risks confusing aspects of the behavioural and cognitive mechanisms involved as well as their function

The scaling pattern we find in humans and some other species appears to be a consequence of the efficiency of internal information flow around networks (West et al. 2020, 2023). In effect, the fractal grouping patterns observed in these species form a harmonic series with very specific values that emerges out of the way individuals organise their personal social relationships. The fact that this pattern is extremely robust (Dunbar 2020) provides us with a formal template by which to judge whether individual species’ social arrangements are genuinely self-organising structures (fractal modularity) or simply casual herd-like gatherings made possible by local ecological conditions (casual modularity).

Here, I compare the modular structure of social groupings in a taxonomically wide sample of mammals using a Horton order analysis (Hamilton et al. 2007; Hill et al. 2008), with the human scaling ratio of 3 as a benchmark. I then ask whether the distinction between those species that exhibit fractal scaling and those that do not lies in their social cognitive competencies. Because data on the social cognitive traits that underpin complex sociality is not widely available outside the primates, I use neocortex ratio as a proxy since this has been shown to correlate with a number of key social cognitive traits in humans and other primates (Dunbar & Shultz 2021a, 2025; Dunbar 2025).

## Methods

I collated data on the mean sizes of different grouping levels for a number of species that have been specifically identified as living in multilevel societies. These include both the primate species listed in the Introduction and a number of non-primate mammals including elephants, orcas, humpback whales and spotted hyaena. Mean grouping sizes for the primates are from Dunbar & Shultz (2021a), except *Rhinopithecus* which is calculated from Qi et al. (2014). Grouping sizes for non-primates are from Dunbar & Shultz (2025), except humpback whales which are from Cantor et al. (2015) (for whom the basal grouping [clique] is taken as mean network degree for individuals).

I use a Horton order analysis (see Hill et al. 2008) to determine whether a species grouping structure is fractal. A Horton order analysis specifies a hierarchically inclusive layering of groupings without defining their sizes, composition or structure. For all the species in the sample, we can specify a basal social group that is substructured into cliques (preferred social partners) (see Dunbar 2025), and I take these to be the first two layers in the Horton order (with the clique defined as order 1, and the basal social group as order 2). All other grouping levels are then naturally assigned to successive orders for as many layers as can be discerned in a species’ data. When group size is plotted on a log_10_ scale, the slope of the regression through the datapoints gives the fractal parameter. Species that have fractal modularity will have a scaling ratio close to *λ=*3.

I use *k*-means cluster analysis of the variance in scaling ratios from *λ*=3 to sort species into distinct clusters for comparison. Finally, I correlate the variance in scaling ratio with species’ neocortex ratio (the ratio of the neocortex volume to the volume of the subcortical brain) where these are available. The neocortex ratio indexes relative investment in neural tissue available for complex cognitive processes compared to managing somatic tissue and basic biological processes like hunger, thermoregulation and sex. For primates, neocortex ratios are from Dunbar & Shultz (2021a) (based on Stephan et al. 1981); those for carnivores and cetaceans are from Dunbar & Shultz (2025).

## Results

Fig. 1 plots the mean layer sizes for the species for which appropriate data are readily available. Notice the tight linearity, and close numerical similarity, in the data for the gelada, hamadryas, chimpanzees and humans, as well as the elephants and orcas. In contrast, the four colobines, Guinea baboon, spotted hyaena and humpback whales exhibit more irregular patterns, as well as possessing only some layers. There is a strong suggestion, for example, that the colobine groupings asymptote at layer 3, whereas the fractal layering in the baboons and humans continues for another two or three layers. In gelada and hamadryas baboons, the upper three Horton order layers correspond to groupings identified as teams, clans and bands; in elephants, they correspond to bond groups, clans and sub-populations (see Hill et al. 2008).

**Fig. 1.**
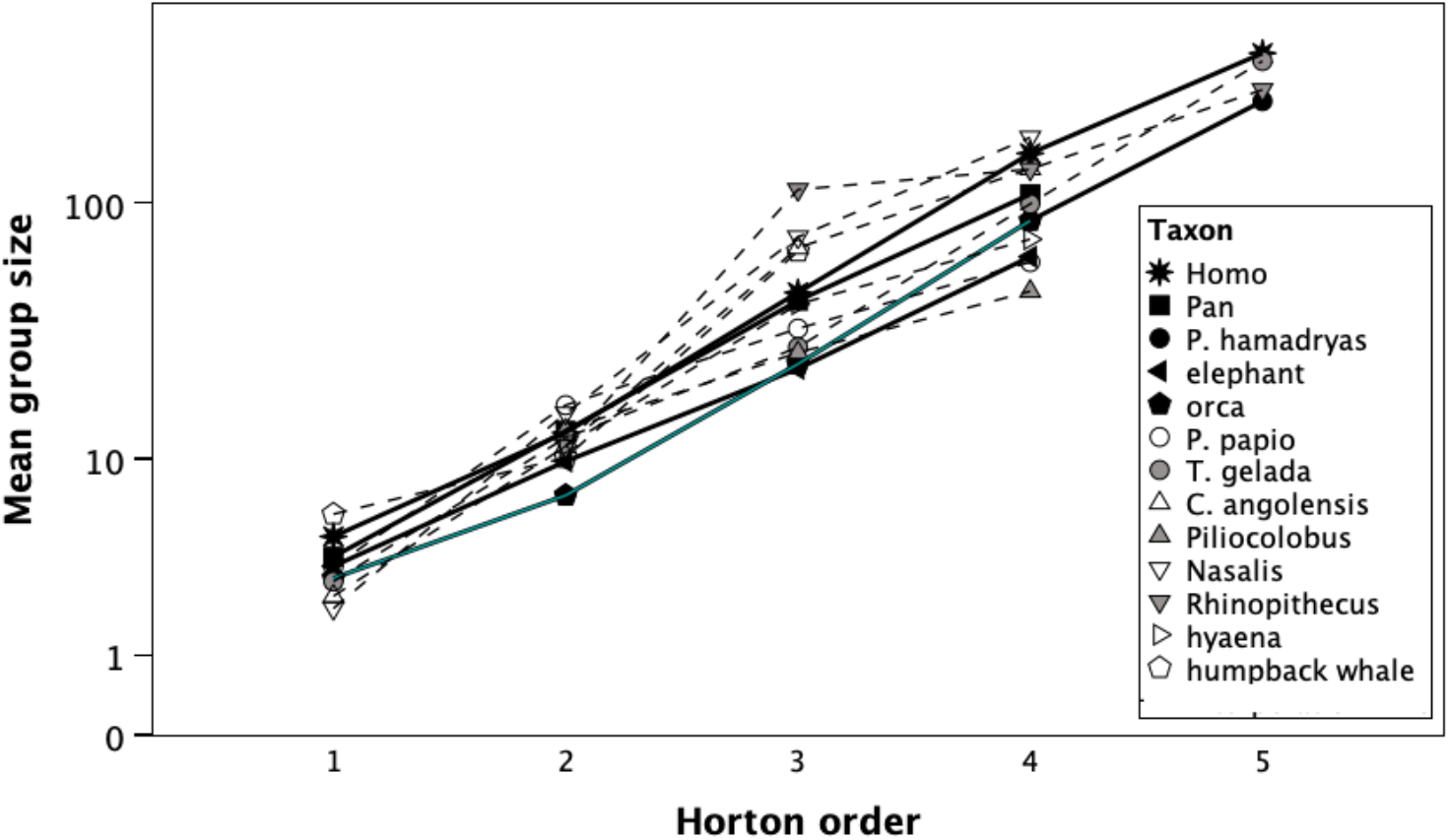
Fractal scaling of group sizes in species that have multilevel social systems. Alliances are grooming cliques (degree plus 1) from Dunbar (2025). Sources: *Crocuta* (spotted hyaena): Dunbar & Shultz (2025); *Colobus angolensis* (Nyungwe population): Stead & Treichot (2019); *Piliocolobus* (red colobus): adapted from Korstjens & Dunbar (2007); *Nasalis* (proboscis monkey): Yaeger (1990, 1991), Matsuda et al. (2024); *Rhinopithecus* (snubnose monkey): Qi et al. (2014, 2017); *Theropithecus gelada* and *Papio hamadryas*: Hill et al. (2008); *P. papio* (Guinea baboon): Fischer et al. (2017); *Pan troglodytes*: Dunbar & Shultz (2021a); orca and elephant: Hill et al. (2008); humpback whale: Cantor et al. (2015); humans: Dunbar (2020). Filled symbols, solid lines: species with formal fractal structures; unfilled symbols, dashed lines: species with casual fractal structures.

Fig. 2 plots the actual scaling ratios for successive grouping levels for each of these species. A *k*-means cluster analysis of the mean variance in the scaling ratios from a predicted value of 3 suggests a partition into two groups whose variances in scaling ratios differ significantly: a low variance group (*Homo, Pan, Papio hamadryas, Theropithecus*, elephants and orcas) and a high variance group (*Papio papio, Colobus angolensis, Piliocolobus, Rhinopithecus, Nasalis*, spotted hyaena and the humpback whale). The first group have highly regular ratios between successive grouping levels, whereas the second group are much more variable (mean log_10_ variance in *λ* = −1.15±0.18SD versus −0.10±0.44; t_11_=5.49, p=0.0002). The social cohesiveness of the 3^rd^ and 4^th^ Horton order groupings of the colobines is often described as weak (Qi et al. 2014; Miller 2019). In this respect, colobine groupings are reminiscent of the spotted hyaena grouping structure: small, coherent breeding units that gather together on an *ad hoc* basis into larger groupings that are inchoately variable in size and organisation (Dunbar 2024).

**Fig. 2.**
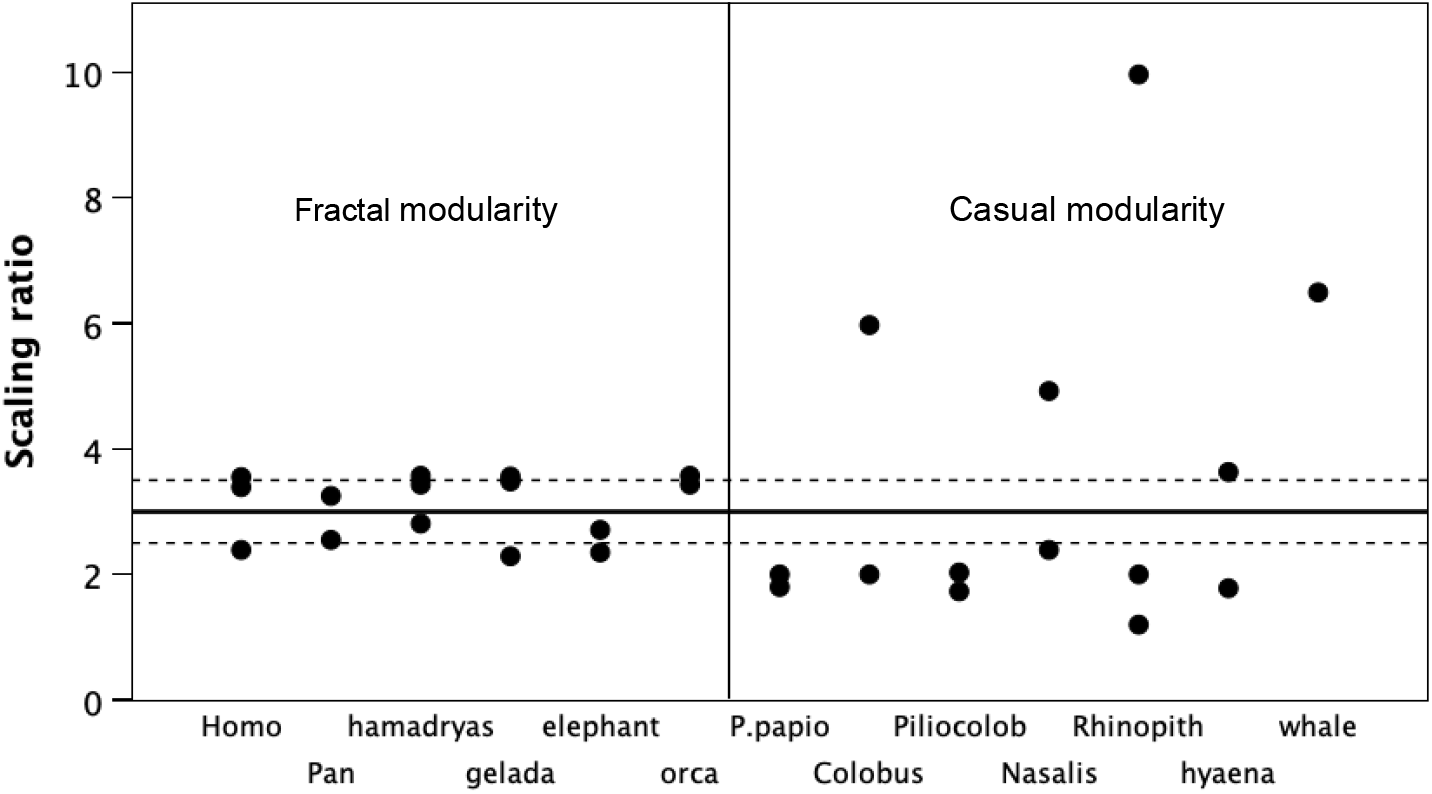
Scaling ratios across grouping layers for the species in Fig. 1. The internal substructure scaling ratio (i.e. that between cliques and basal social units) is not included. The thick horizontal line indicates the scaling ratio of 3 characteristic of a fractally structured social system; dashed lines indicate range 2.5-3.5. Vertical line differentiates fractal from casually modular species yielded by a *k*-means cluster analysis of the species’ mean variances. Piliocolob = *Piliocolobus*; Rhinopith = *Rhinopithecus*; whale = humpback whale.

To test whether species with fractal modularity invest more heavily in cognition than in managing somatic tissue, I plot neocortex ratio for the two clusters in Fig. 3. Fractal species have significantly larger neocortex ratios (t_8_=-2.47, p=0.039). This suggests that there is likely to be a difference in the kinds of cognitive skills (and, in particular, social cognitive skills) possessed by the two groups of species.

**Fig. 3.**
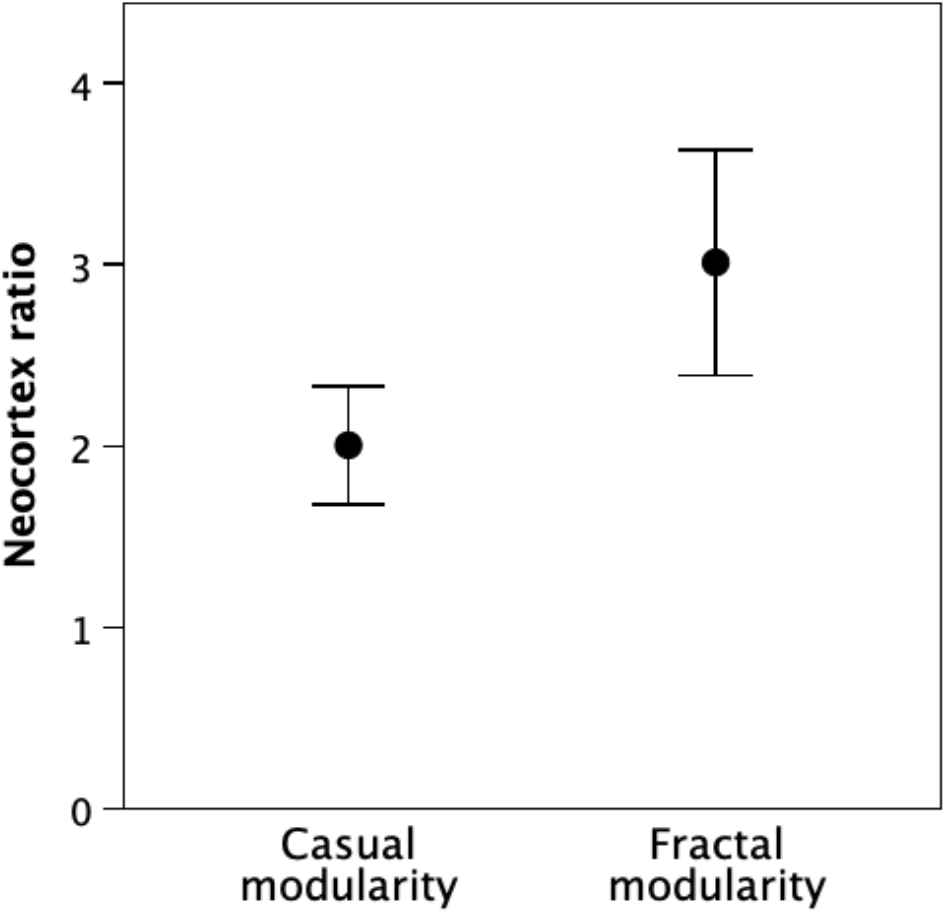
Mean (±2 sem) neocortex ratio for species in Fig. 2 that have casual versus fractal modularity. Sample sizes are N=5 and N=6, respectively.

## Discussion

Although many species have been described as having multilevel grouping patterns, analysis of the fractal structure of these groupings suggests that two different kinds of social style can be differentiated: casual modularity and fractal modularity. Species that have a casually modular form of sociality are in many ways more similar to herding ungulates: basal social units gather together on an *ad hoc* basis into larger groupings that lack cohesiveness, are structurally variable and, in all probability, do not involve personalised knowledge of other members. In these cases, scaling ratios in the basal layers approximate those of fractally modular species, but those in the upper layers are more variable and uncorrelated with those in the basal layers. In contrast, species with a fractally modular form of sociality seem to create higher level groupings by bolting together lower level groups on what is essentially an all-or-none basis, and do so by exploiting the same mechanisms used in bonding basal social units (see Dunbar 2025). Hence, in this case, the scaling ratios are remarkably consistent across grouping levels.

This suggests that an important distinction needs to be drawn between those species that exhibit fractal modularity (where successive layers are built up by bolting together a set number of groups from a lower level) and those that have more of a herd-like structure (i.e. large groups that gather casually, for example on rich food resources, but have no formal structure) (casual modularity). Notice that the same fractal structure is found both in species that exhibit individual-level fission-fusion sociality (chimpanzees, elephants and dolphins, and perhaps humans) and in species that have group-level forms of fission-fusion sociality (hamadryas and gelada baboons, where stable harems join together in herds), as well as in species that have male philopatry (chimpanzees, and maybe humans) and those that have female philopatry (hamadryas and gelada). In other words, the distinction between fractal and casual modularity cuts across more conventional distinctions between different forms of sociality.

Since large neocortex ratios correlate with greater abilities on a number of cognitive skills (including mentalising and inhibition) that, in primates, are crucial for living in large stable groups (Dunbar & Shultz 2021a, 2025; Dunbar 2025), the fact that fractally modular species have larger neocortex ratios than casually modular species (Fig. 3) suggests that all these species, irrespective of taxonomic order, may be able to draw on more sophisticated forms of cognition. What seems to lie at the heart of fractal modularity, then, is the capacity to deploy novel cognitive abilities that allow individuals to manage relationships with individuals with whom they do not necessarily live on a daily basis. These cognitive strategies allow them to extend the effective size of their groups by several more layers by deferring group fission in a way that creates a large substructured grouping. This contrasts with the smaller-brained colobines that have a small fixed group size, but can fuse these temporarily in groupings (herds) of variable size that depend on who happens to be around. This is signalled in Fig. 1 by the fact that the three of the four colobines have significantly larger groupings at level 3 than the papionines and humans (t_4_=2.67, p=0.056) but not at level 2 (t_4_=0.48, p=0.654). Their level 3 groupings are not modular fractals of a lower level grouping. Like herding in ungulates, casual modularity does not require individuals to recognise each other or to have stable relationships. Instead, very simple rules such as “always submit to a more powerful individual” are all that is needed to regulate relationships. The distinction is similar to that drawn by Lukas & Clutton-Brock (2018) between species that have rule-based relationships (structural or organisational complexity) and species that create stable relationships (relational complexity).

It is noteworthy that all the fractal species occupy habitats (terrestrial or oceanic) with high predation risk and few available refuges compared to those occupied by most of the casually modular species. Baboons that live in habitats with few trees (forest cover <25% of the ground surface) and high predator densities (>0.25 individual predators/km^2^) live in larger groups than those that live in more heavily forested habitats (Dunbar & MacCarron 2019). Similarly, dolphins (but not humpback whales) face a considerable predation threat from sharks (Heithaus 2001).

Casually modular species can also face significant predation risk, and this may account for the fact that some, but not all, of their populations sometimes live in large groups. The Nyungwe colobus, for example, suffer considerable predation from chimpanzees (Miller 2019), as do some red colobus (*Piliocolobus* spp.) populations (Stanford 1998; Surbeck & Hohmann 2008; Watts & Amsler 2013). *Piliocolobus* populations that are sympatric with chimpanzees live in much larger groups (mean 46.0, range 23-60) than populations of this genus from areas where chimpanzees do not occur (mean=22.1, range 13-27) (Korstjens & Dunbar 2007). This suggests that while herd formation may be a natural response to locally high predation risk, only when this is characteristic of all the habitats typically occupied by a species is the selection pressure sufficiently consistent to promote the evolution of fractal modularity.

Most of the species sampled are capable of living in large groups under more benign ecological conditions. Time budget models for *Colobus, Papio* and *Theropithecus* all indicate that these genera have considerable ecological flexibility. As a result, large group sizes are possible in ecologically more benign habitats if and when large groups are locally advantageous. The black-and-white colobus model, for example, predicts that limiting group size will be as high as 120 in habitats where rainfall is around 2900mm and mean temperature exceeds 30°C (Korstjens & Dunbar 2007). *Papio* group sizes might be as large as 400 under similar conditions (Dunbar 1992a) – as is typically the case in the West African equatorial forest habitats where *Mandrillus* habitually occurs in groups of this size. At altitudes of 3000-3500m asl where grasses are more succulent and less silicated, gelada can – and do – live in herds of 200-400, whereas at lower altitudes herd sizes are much smaller (individual harems of 10-15 animals typically forage alone at altitudes below 1000m asl: Dunbar 1992b).

In fact, many, though not all, of the colobine populations with very large groups occupy high altitude habitats (>1000m asl) that are cool (mean ambient temperatures 5-15°C) and wet (annual rainfall >750mm) (Nyungwe and Ruwenzori *Colobus, Rhinopithecus*), heavily forested (Nyungwe and Ruwenzori *Colobus, Nasalis, Rhinopithecus*), often poor quality habitats (e.g. Nyungwe *Colobus* and *Rhinopithecus* diets are heavily based on lichens). Cooler, wetter habitats are in general characterised by forage that is more digestible, and this may be especially important for colobines that are dependent on leaves that require considerable time for fermentation. Fimbel et al. (2001) noted that the Nyungwe habitat was characterised by high protein:fibre ratios, which will reduce the time needed for fermentation. As a result, the Nyungwe *Colobus* and *Rhinopithecus* populations devote significantly less time to resting than do other *Colobus* populations: Nyungwe: 32% (Fashing et al. 2007); *Rhinopithecus*: 28.3%±8.2 sem, N=6 sites (Ding & Zhao 2004; Guo et al. 2007; Xiang et al. 2010; Liu et al. 2013; Li et al. 2015, 2023); all other *Colobus:* 58.5±8.3%, N=10 sites (Korstjens & Dunbar 2007) (t_15_=7.49, p<<0.0001).

Living in large groups, however, creates very significant stresses that dramatically impact female fertility (the ‘infertility trap’: Dunbar & Shultz 2021b). Females need to find ways to minimise these stresses. I suggest that we can interpret the distinction between fractally and casually modular species as alternative solutions to this problem. One solution is to adopt some form of herding: individuals join a herd when it is advantageous to do so, but leave as soon as the costs exceed the benefit. This requires little more than simple joiner-leaver principles. Moreover, since relationships in herds are generally anonymous and rarely involve repeated meetings with the same individuals over time, only the very simplest rules of engagement are necessary (‘always submit to bigger individuals’). As a result, no new forms of cognition are required. The alternative is to form alliances that will buffer individuals against the stresses, thus allowing females to remain together in larger groups. Such a strategy comes with a cost, because it involves creating bonded relationships that can act as stable coalitions. This creates gravitational forces that make it difficult for individuals to leave groups on their own (Dunbar 2025).

More importantly, creating effective, stable alliances is expensive both because of the time needed to build and maintain stable relationships and because the cognitive skills needed to manage relationships of different kinds require investment in novel, neurally expensive brain circuits (Dunbar 2025). There is considerable evidence for novel forms of social cognition from many of the taxa in the fractal species and their allies, including understanding third party relationships, reconciliation (repairing weakened relationships), behavioural inhibition (self-control), the capacity to assess individual on two dimensions simultaneously (e.g. kinship versus dominance) and mentalising (understanding others’ intentions), with little evidence for these in the group of casually modular species and their allies (Dunbar & Shultz 2021a, 2025; Dunbar 2025). There is urgent need for better informed studies of social cognition in a wider range of primate and non-primate species to allow these cognitive differences and their demographic implications to be explored in more detail.

In sum, these results thus suggest that there may be two routes to creating superlarge multilevel societies. One is a bottom-up process that allows individuals or small harem units to coalesce into loose herds when they need to; the other is a top down process that arises when larger groups can be substructured into fractally organised subunits. I suggest that the former requires no new forms of cognition, whereas the other requires the evolution of novel cognitive abilities that a neurally expensive.

